# The Cryptic Bacterial Microproteome

**DOI:** 10.1101/2024.02.17.580829

**Authors:** Igor Fesenko, Harutyun Sahakyan, Svetlana A. Shabalina, Eugene V. Koonin

## Abstract

Microproteins encoded by small open reading frames (smORFs) comprise the “dark matter” of proteomes. Although functional microproteins were identified in diverse organisms from all three domains of life, bacterial smORFs remain poorly characterized. In this comprehensive study of intergenic smORFs (ismORFs, 15–70 codons) in 5,668 bacterial genomes of the family Enterobacteriaceae, we identified 67,297 clusters of ismORFs subject to purifying selection. The ismORFs mainly code for hydrophobic, potentially transmembrane, unstructured, or minimally structured microproteins. Using AlphaFold Multimer, we predicted interactions of some of the predicted microproteins encoded by transcribed ismORFs with proteins encoded by neighboring genes, revealing the potential of microproteins to regulate the activity of various proteins, particularly, under stress. We compiled a catalog of predicted microprotein families with different levels of evidence from synteny analysis, structure prediction, and transcription and translation data. This study offers a resource for investigation of biological functions of microproteins.

**Highlights:**

- Thousands of previously unknown bacterial microproteins predicted
- Most microproteins belong to lineage-specific families, revealing unexplored diversity of bacterial proteomes
- Comparative genome analysis suggests de novo emergence of numerous microproteins
- Interactions between stress-induced microproteins and known functional proteins predicted
- This study provides a resource to investigate cryptic bacterial microproteomes

## Introduction

Small open reading frames (smORFs), usually defined as ORFs shorter than 100 codons in eukaryotes^1^ and shorter than 50 or 70 codons in prokaryotes^2^, remain poorly annotated and barely investigated due to the limitations in gene prediction by computational methods and thus comprise part of the “dark matter” in all genomes^3^. Although translated smORFs are found in various genomic locations in all organisms, the biological functions (if any) of the encoded microproteins are not known, and therefore they are hard to distinguish from translational noise^4^. Nevertheless, some microproteins have been functionally characterized in different models, including animals, plants, fungi, and bacteria^5–9^. Moreover, a standardized catalog of more than 7,000 human smORFs validated with Ribo-seq ORFs was built in a recent community effort^10^, emphasizing the growing importance of the microproteome.

Microbiome analysis led to the identification of tens of thousands of small proteins in bacteria^11,12^. In prokaryotic genomes, intergenic smORFs (ismORFs) could comprise an important source of new proteins emerging from non-coding sequences and potentially involved in a variety of cellular functions^13,14^, but to our knowledge, systematic analysis of prokaryotic ismORFs has not been reported^2^.

*De novo* gene birth is thought to play a major role in the emergence of microproteins, and most smORF-encoded microproteins appear to be evolutionarily young^15^. The processes of *de novo* gene birth have been studied in a range of species, such as human^16^, mouse^17^, yeast^18^ and plants^19^. The properties of such young microproteins are almost indistinguishable from those of random sequences, and most of the microproteins appear to be highly disordered^20,21^. These studies notwithstanding, there is no comprehensive understanding of the extent and impact of de novo gene birth and death processes on the microproteome dynamics, especially in bacteria. We performed a comprehensive analysis of millions of ismORFs longer than 15 aa in 5668 bacterial genomes of the family Enterobacteriaceae and characterized the cryptic bacterial microproteome. Through analysis of evolutionary conservation, we predicted tens of thousands of microprotein families, most of which are hydrophobic and have low sequence complexity. We predict that stress-regulated microproteins interact with larger, functionally characterized proteins, suggesting that microproteins play important roles in stress response.

## Results

### Intergenic sequences (IGs) as the birthplace of genes

#### Intergenic sequences are short and AT-rich

The evolutionarily young microproteins encoded by small open reading frames (smORFs) that often evolve *de novo*^15,22^ remain largely unidentified in bacterial genomes. Therefore, considering intergenic regions as a reservoir of unidentified small proteins and the potential birthplace of new genes, we extracted intergenic sequences (IGs) from 5668 bacterial genomes belonging to 23 bacterial genera in the family *Enterobacteriaceae*. Next, 17,056,700 IGs with lengths ranging from 33 to 1500 nt were selected for further analysis (Figure 1A). Within all genera, the mean length of the selected IGs was about 200 nt, with an interquartile range (IQR) of 175-212 nt (Figure 1B). Thus, as expected, bacterial IGs are long enough to accommodate numerous microproteins but not larger proteins non-overlapping with annotated protein-coding genes.

**Figure 1.**
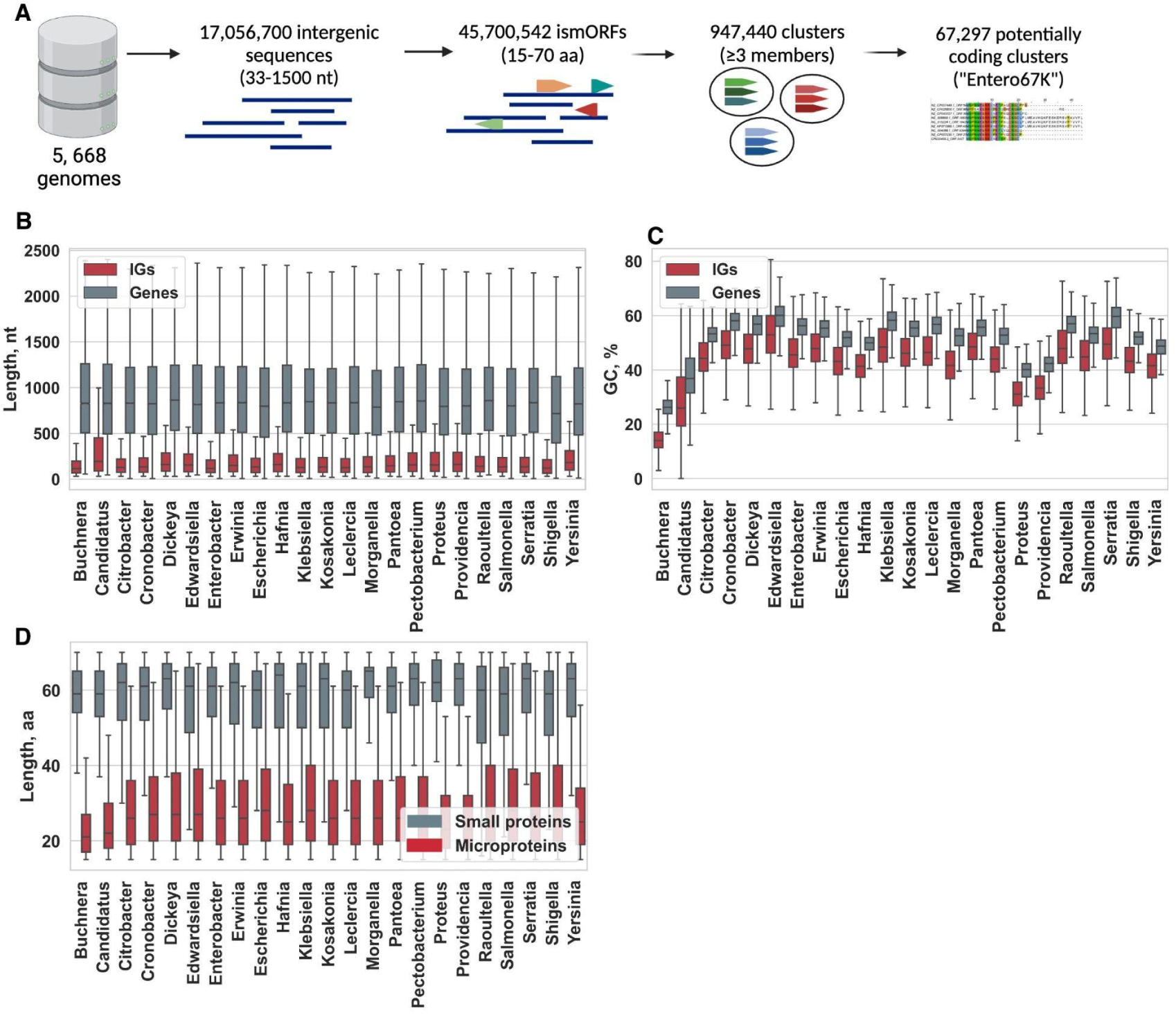
Intergenic sequences and and intergenic small ORFs in enterobacteria. **(A)** Pipeline for identification and analysis of intergenic small Open Reading Frames (ismORFs). More than 17 million intergenic sequences from 23 bacterial genera in the family *Enterobacteriaceae* were used to predict intergenic smORFs. Predicted microproteins were clustered by the CD-HIT tool^25^ in 947,440 clusters with a median length of 15 to 70 aa, and at least three sequences were selected for further analysis. Signatures of evolutionary selections were used to define the set of potentially coding clusters (Entero67K); **(B)** The lengths of the selected intergenic sequences and genes from RefSeq annotation across all genera; **(C)** The GC-content of annotated genes and intergenic sequences across all genera; **(D)** The lengths of annotated small proteins and putative microporteins encoded by intergenic smORFs across all genera (Mann-Whitney U test, *P* < 10^-10^ for all comparisons).

The GC content of coding sequences in 23 genera of enterobacteriaceae ranged from 26.3% (in *Buchnera* spp.) to 57.2% (in *Klebsiella* spp.). The AT content in IGs was consistently and significantly higher than in coding sequences across all genera (Mann-Whitney *U* test, *P* < 10^-15^ for all comparisons; Figure 1C). The base composition of IGs can partly shape the properties of emerging microproteins. Given the structure of the genetic code, these sequences would favor the formation of hydrophobic microproteins.

#### Prediction of small ORFs in intergenic regions

The size limit for including a protein in the RefSeq database (National Center for Biotechnology Information, NIH) based on *ab initio* prediction alone is 40 aa^23^. Given that even some bacterial microproteins as small as 15 amino acids have been demonstrated to be functional^24^, we identified a set of 45,700,542 bacterial ismORFs between 15 and 70 codons in length, beginning with the start codons AUG, GUG, or UUG. We refer to the hypothetical proteins encoded by these ismORFs as microproteins by analogy with the eukaryotic smORF-encoded microproteins^5,15^.

The median length of putative microproteins encoded by ismORFs was 27 aa (IQR = 26.5-28 aa) across all genera. As a control, we extracted from RefSeq a set of 562,920 small genes encoding annotated small proteins ≤ 70 aa, excluding genes annotated as “putative” or “uncharacterized” that might represent erroneously predicted protein-coding genes, from the corresponding bacterial species. The small proteins were, on average, significantly longer than the putative microproteins across all genera (Figure 1D).

### Identification of families of predicted microproteins

Previous studies^13,26^ suggest that genomes encompass different types of potentially translated microproteins, including a continuum of likely protogenes^4^. To distinguish random ismORFs from potentially coding ones, we focused on the estimation of evolutionary selection signatures as a signature of functional microproteins. A number of methods for differentiation of sequences with coding potential from non-coding regions have been proposed, mainly based on experimental data^27,28^ or sequence features of already annotated small proteins in certain species^29,30^. For further analysis, ismORFs-encoded putative microproteins were clustered using the following parameters: at least 50% amino acid identity in an alignment spanning at least 95% of the microprotein sequences. This procedure resulted in 947,440 microprotein clusters, including at least three sequences each, with median lengths of 15 to 70 aa (Figure 1A). We also clustered 562,920 annotated functional small enterobacterial proteins (≤ 70 aa), encoded by the set of small genes defined above, into 6107 genus-specific clusters (hereafter “SmallProt”; Table S1).

#### Coding potential of ismORFs

To predict which ismORFs encoded putative microproteins in enterobacterial genomes, two computational methods were employed, RNAcode tool^31,32^ and modified EvolScore^5^, both based on the estimation of selective constraints (see STAR Methods for details). RNAcode was recently used to identify functional small proteins in metagenomic analysis^11,33^. Using the RNAcode tool (*P* < 0.05), we classified about 3% of microprotein clusters as “coding” (Figure 2A and B). However, because only ∼30% of SmallProt clusters met the criteria to be considered “coding” by RNAcode, this tool clearly has a high false negative rate. Therefore, to identify potentially coding ismORFs, we additionally developed the EvolScore approach, which is based on calculations of evolutionary scores: *dN* (non-synonymous substitutions), *dS* (synonymous substitutions), *dN/dS* ratio, and p-distances (*Kd*; Kimura two-parametric model, K2P). Because evolutionary rates of small proteins in *Enterobacteria* are generally unknown^34^, to select reliable thresholds for EvolScore, we calculated extreme quantiles (1% or 99%), for *dN*, *dS* and *Kd* values in SmallProt clusters and shuffled microprotein clusters with the bootstrap (see STAR Methods for details; Figure 2C).

**Figure 2.**
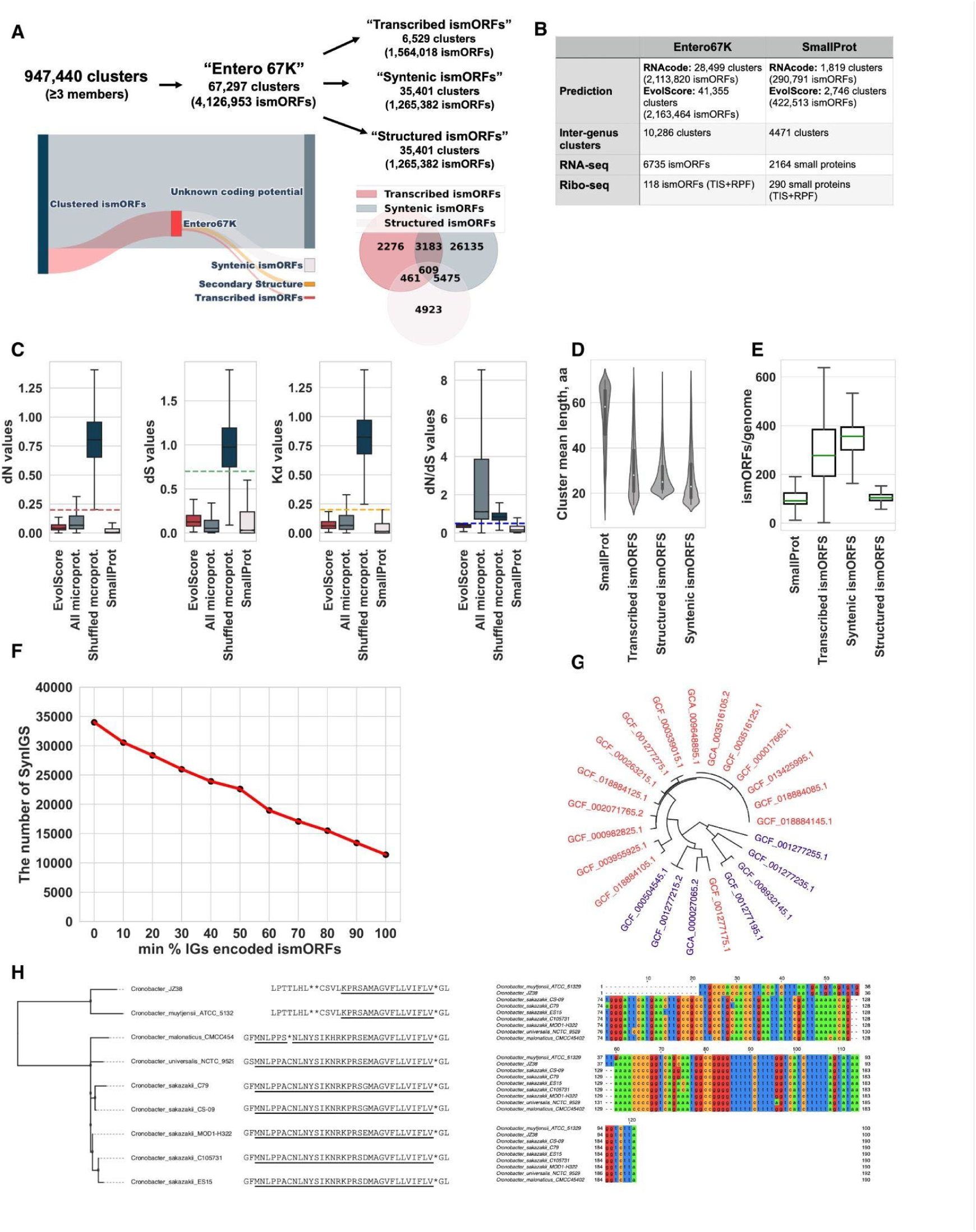
Families of ismORF-encoded predicted microproteins. **(A)** Sankey plot showing the proportion of clustered ismORFs and those with that were predicted as coding. The microprotein families have evidence of transcription (“Transcribed ismORFs”, 15 species), located in syntenic intergenic regions (“Syntenic ismORFs”), and have predicted structure (“Structured ismORFs”). **(B)** RNAcode^32^ and MegaX^36^ tools were used for the prediction of the coding potential of intergenic smORFs. Based on this analysis, the “Entero67K” set of potentially coding microproteins (15–70 aa) was established. **(C)** The calculated evolutionary rates for ismORFs classified as “coding” (“EvolScore”), all clustered ismORFs, shuffled microprotein clusters, and annotated small proteins (SmallProt); EvolScore thresholds are shown by dashed lines; **(D)** The length of annotated small proteins in the “SmallProt” dataset and ismORFs from “Entero67K” families; **(E)** A boxplot showing the number of ismORFs from Entero67K families in enterobacterial genomes; **(F)** The plot shows the number of Syn-IGS blocks (axis Y) in relation to a minimal proportion of genomes (axis X) containing an intact ismORF (Entero67K) in the corresponding Syn-IGS; The plot was built for intergenic sequences, containing ismORFs from one Entero67K cluster; **(G)** The phylogenomic tree of the genus *Cronobacter*. The species with intact ismORFs from the syntenic cluster “Cronobacter_15372” are shown in red; **(H)** Non-redundant nucleotide and translated sequences of the corresponding Syn-IGS (for the cluster “Cronobacter_15372”) are shown.

Applying the two approaches to the SmallProt dataset resulted in ∼30% and ∼45% clusters classified as “coding” by RNAcode and EvolScore, respectively, with a limited overlap between the two sets of predictions (Figure S1. Related to Figure 2), suggesting that there is no universal solution for predicting the coding potential of short ORFs. Thus, prediction using different methods for the detection of selection, analysis of genomic synteny, and published experimental data (see below), need to be combined for a comprehensive search for microprotein candidates.

To further validate the microprotein prediction, the random forest (RF) algorithm was trained on a randomly chosen set of SmallProt clusters as the positive dataset and randomly chosen shuffled microprotein clusters as the negative dataset. The model was evaluated with respect to precision, recall, and Area Under the Receiver Operating Characteristics (AUROC) values. The AUROC value for the model was 0.961. The evaluation parameter precision (defined as the proportion of true positives among all predicted positives) was 0.948, with recall (sensitivity) reaching 0.90 (Figure S1. Related to Figure 2). Thus, the model was found to be both highly sensitive and highly specific. Using this model, we predicted 4.3% of the microprotein clusters as “coding” (Figure 2A and B). The evolutionary rates of these clusters of predicted microproteins were comparable to those of the annotated small proteins (Figure 2C). The set of ismORFs predicted as coding using the RF model and the set of predictions obtained using the EvolScore thresholds overlapped almost completely (>99%), asserting the robustness of the prediction.

The union of the microprotein clusters identified as coding with at least one approach (RNAcode and/or EvolScore) is hereafter referred to as the Entero67K microprotein set (Figure 2A; Table S2). The Entero67K set of predicted microproteins is notably larger and possesses shorter sequences (Figure 2D; Kruskal-Wallis rank sum test, *P* < 10^−15^) compared to the set of annotated small proteins (SmallProt; Figure 2E). Approximately 51% of the Entero67K clusters are highly conserved and show strong signatures of purifying selection, where *Kd*<0.1 and *dN/dS*<1. However, a substantial number of microproteins had *dN/dS*>1, suggesting that they might be subject to positive selection (see below). Thus, our findings suggest that the bacterial microproteome remains largely uncharacterized and contains microproteins of different evolutionary ages.

#### Synteny conservation in ismORF families

We next explored how many ismORFs from the Entero67K set occupied the same position in syntenic regions of enterobacterial genomes. To this end, we identified syntenic regions in the genomes of each enterobacterial genus using the SibelliaZ tool^35^, which compares nucleotide sequences of multiple genomes and uses a compacted de Bruijn graph to generate locally-collinear blocks (LCBs). Altogether, 3,095,516 LCBs were identified, with genome coverage (fraction of the genomic sequence included in the alignment) ranging from 0.75 for *Buchnera* to 0.98 for *Pantoea*.

We next identified syntenic intergenic regions based on the intersection between LCBs and intergenic sequences (see Methods). Intergenic regions were combined into one syntenic block (Syn-IGS) if 1) they intersected the same LCB by at least 50%; 2) intergenic sequences from at least two different species were represented; and 3) at least one intergenic sequence contained an ismORF from the Entero67K set. This analysis produced 38,759 Syn-IGSs that included 2,985,292 intergenic sequences.

The Syn-IGSs, on average, contained 1.4 microprotein clusters, indicating that in most cases, a single smORF with predicted coding potential was found within an intergenic region. Overall, ∼30% of ismORFs (Entero67K) were found to be encoded in syntenic intergenic regions (“Syntenic ismORFs”; Figure 2A). This relatively low level of synteny conservation could be caused by different reasons, including the methodological constraints of synteny analysis by SibelliaZ or by actual genomic rearrangement in IGs that results in short LCBs that do not meet our filtration criteria.

We next roughly estimate the turnover rate of ismORFs based on their presence or absence in Syn-IGSs. In ∼34% of the Syn-IGSs, each intergenic sequence (from one syntenic block) was found to contain an intact ismORF from the same Entero67K cluster (Figure 2E). Thus, a majority of the syntenic blocks contain intergenic sequences lacking intact ismORFs suggesting that many ismORFs evolved from non-coding sequences relatively recently, after the divergence of the respective bacterial species, or have been lost in some species. An example of a synteny intergenic block is shown in Figures 2F and G. Taken together, our results demonstrate a high turnover rate of ismORFs in intergenic regions, suggesting that lineage-specific emergence of *de novo* genes is common in the evolution of enterobacteria.

### Evidence of transcription and translation of ismORFs

#### Transcription of ismORFs

Next, we searched for evidence of transcription of ismORFs. To this end, 71 publicly available RNA-seq transcriptomes of enterobacterial species from 15 genera were analyzed (Table S3). On average, about 50% of all ismORFs had a transcriptional level of >1 TPM (transcripts per million) and were fully covered by reads. Overall, we confirmed transcription of 59,402 genes and 6,735 ismORFs (median=449 transcribed ismORFs/genome). On average, 3960 genes/genome, including 144 small genes/genome (SmallProt dataset), were transcribed with TPM > 1 (Table S3). However, our estimations of transcribed genes and ismORFs might be overestimated due to the absence of strand-specific libraries for most species.

As expected, the transcription level of ismORFs (median=16 TPM; interquartile range (IQR) = 5.04 - 56.6) was significantly lower than that of annotated genes (median=31.6 TPM; IQR=8.07-101.5) in all genera (Mann-Whitney U test, *P* < 10^-7^). Thus, intergenic ismORFs appear to be extensively transcribed, even at comparatively low levels, facilitating *de novo* gene birth. The set of transcribed ismORFs from 15 bacterial genera with signatures of evolutionary selection (that is, belonging to Entero67K), is hereafter referred to as “MicroProt6K” (Figures 2A and B).

We further estimated the proportion of ismORFs that shared a common primary transcription start site (pTSS) with nearby genes. Remapping “Cappable-seq” data for the *E. coli* K-12 str. MG1655 (”NC_000913.3” genome; “Ettwiller set”) strain, we identified 16,587 potential TSSs (Table S3), which were closely similar to those previously reported^37^. There was a significant difference (Kruskal-Wallis rank sum test, *P* < 10^−15^) between the distances to the nearest pTSS for annotated genes (median = 119 nt) and ismORFs (median = 546 nt) suggesting that most of the ismORFs are not associated with typical pTSS. However, we found that, of the 661 Entero 67K ismORFs represented in the NC_000913.3 genome, 114 ismORFs shared a common pTSS with downstream genes, and 97 shared a pTSS with the upstream gene. Moreover, 32 ismORFs shared a common pTSS with both the upstream and downstream genes, suggesting that they are parts of the respective operons.

#### Translation of ismORFs

To identify translated ismORFs in the genomes of *E. coli* and *Salmonella enterica*, we searched the data from ribosome profiling experiments (Ribo-seq; Table S4). The data from both types of Ribo-seq experiments, mapping translation initiation sites (TISs)^38,39^ and ribosome protected fragment data (RPFs)^40–42^, were searched. To identify potentially translated ismORFs, the smORFer tool^28^, which outperforms other tools in the detection of bacterial smORFs^40^, was used.

The analysis of TIS experiments results with Onc112 (*E. coli* MG1655)^39^ and retapamulin (*E. coli* MG1655 and BL21 stains)^38^, combined with RPFs data, revealed putative translation of ∼67% (108/160) SmallProt ORFs in *E. coli* MG1655 and ∼57% (82/143) SmallProt ORFs in *E. coli* BL21 strains (Table S4). Over 90% of these potentially translated small proteins start at an AUG codon (Figure 3A). In addition to the small proteins, 364 ismORFs in *E. coli* MG1655 and 765 ismORFs in *S. enterica* exhibited both TIS and RPF signals. The TIS signals of potentially translated ismORFs were significantly lower compared to the small proteins (Mann-Whitney U test, *P* < 10^-15^), suggesting that many of these might result from spurious translation or that the respective ismORFs are translated at a low level in normal conditions, in agreement with previously demonstrated trends^4,39^. However, ismORFs with a TIS signal comparable to annotated small proteins were also identified (see example in Figure 3B). Overall, Ribo-seq TIS data combined with traditional RPFs data showed evidence of translation for ∼62% of the annotated SmallProt ORFs, but only ∼6.5% of the ismORFs in both *E. coli* strains (Table S4).

**Figure 3.**
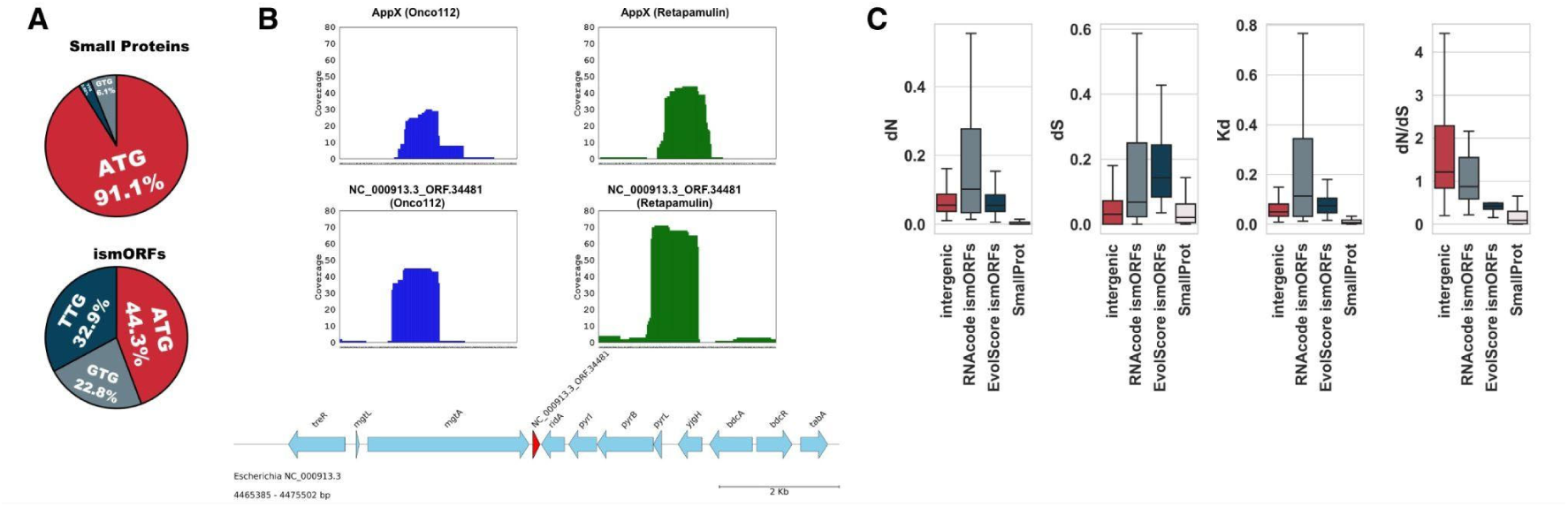
Analysis of potentially translated ismORFs. **(A)** Proportion of “ATG”, “GTG”, and “TTG” start codons in translatable small proteins and Entero67K ismORFs; **(B)** The comparison of TIS read coverage for the known small protein AppX and microprotein NC_000913.3_ORF.34481. The microprotein has signals from both TIS and RPF data and is located downstream of the protein-coding gene MgtA. In *Salmonella spp.,* two small proteins, MgtR and MgtU, interacting with MgtA, are located in this position; **(C)** The evolutionary rates of potentially translated ismORFs without predicted coding potential (“intergenic”), ismORFs from the Entero67K set, predicted by different methods (RNAcode and EvolScore) and small proteins (SmallProt). The evolutionary rates significantly differ: *dN* - Kruskal-Wallis rank sum test, *P* < 0.001; *dS* - Kruskal-Wallis rank sum test, *P* < 10^-15^; *Kd* - Kruskal-Wallis rank sum test, *P* < 10^−5^; *dN/dS* - Kruskal-Wallis rank sum test, *P* < 10^−15^.

We additionally examined traditional ribosome profiling data from *S. enterica* serovar Typhimurium str. SL1344 and 14028S and *E. coli* O157:H7 Sakai. The analysis of Ribo-seq data from the *E. coli* O157:H7 Sakai yielded additional 168 ismORFs from Entero67K, each with at least 70% RPF read coverage (Table S4). In addition, 227 ismORFs in *S. enterica* str. 14028S and 149 ismORFs in *S. enterica* str. SL1344 also belonged to Entero67K (Table S4). Altogether, about 10% of potentially translated ismORFs belonged to the Entero67K set, that is, showed the signature of purifying selection. Thus, the substantial majority of potentially translated ismORFs showed no signs of selection. Because the *dN/dS* values of potentially translated ismORFs were highly variable (median = 1.53; IQR = 0.84–7.6; Figure 3C), we analyzed the signatures of positive selection in the set of all potentially translated ismORFs using the HyPhy BUSTED algorithm^43^. This method tests if a gene experienced positive selection at at least one site on at least one branch of the respective phylogenetic tree. The fraction of potentially translated ismORFs that displayed evidence of episodic positive selection (24%) was statistically indistinguishable from the proportion of small proteins (SmallProt) containing at least one positively selected site (HyPhy BUSTED; *P*adj < 0.05). These findings suggest that episodic positive selection is common among ismORFs and similar to the extent of this type of selection in annotated genes encoding small proteins.

### Characteristics of microprotein families

#### Evolutionary conservation of predicted microproteins

The majority of functional bacterial small proteins are conserved in only a limited number of organisms^44^. Moreover, identification of orthologous small proteins is complicated by the short length of the coding sequences, the fact that many small proteins are hydrophobic and can show spurious sequence similarity, and the non-uniform annotation of small protein genes in completed genome sequences^44^. We next compared the conservation of small protein (SmallProt) and microprotein families (Entero 67K) using sensitive profile-profile search by the HHsuite tool^45^.

In the SmallProt dataset, for 73% of the small protein clusters, at least one homologous cluster was identified in other genera (Figure 2B; Table S5). However, only 0.2% of SmallProt clusters were conserved across the entire set of the analyzed enterobacterial genomes (Figure 4A and B). In contrast to annotated small proteins, only about 13% of the Entero67K clusters had at least one homolog in other genera (Figure 2B; Table S5). Among these conserved clusters, ∼53% included homologs from more than two genera, as opposed to 85% in the SmallProt clusters (Figure 4A), indicating that the vast majority of the microproteins are lineage-specific. Most of the conserved clusters included predicted microproteins from the genera *Citrobacter* and *Klebsiella*; *Escherichia* and *Salmonella; Klebsiella* and *Pectobacterium* (Figure 4C). Analysis of the amino acid alignments of the microproteins from different genera revealed pronounced differences in sequence lengths (mean difference between sequences, 13 aa), suggesting frequent sequence changes beyond the conserved core of the microprotein clusters (see example in Figure 4D).

**Figure 4.**
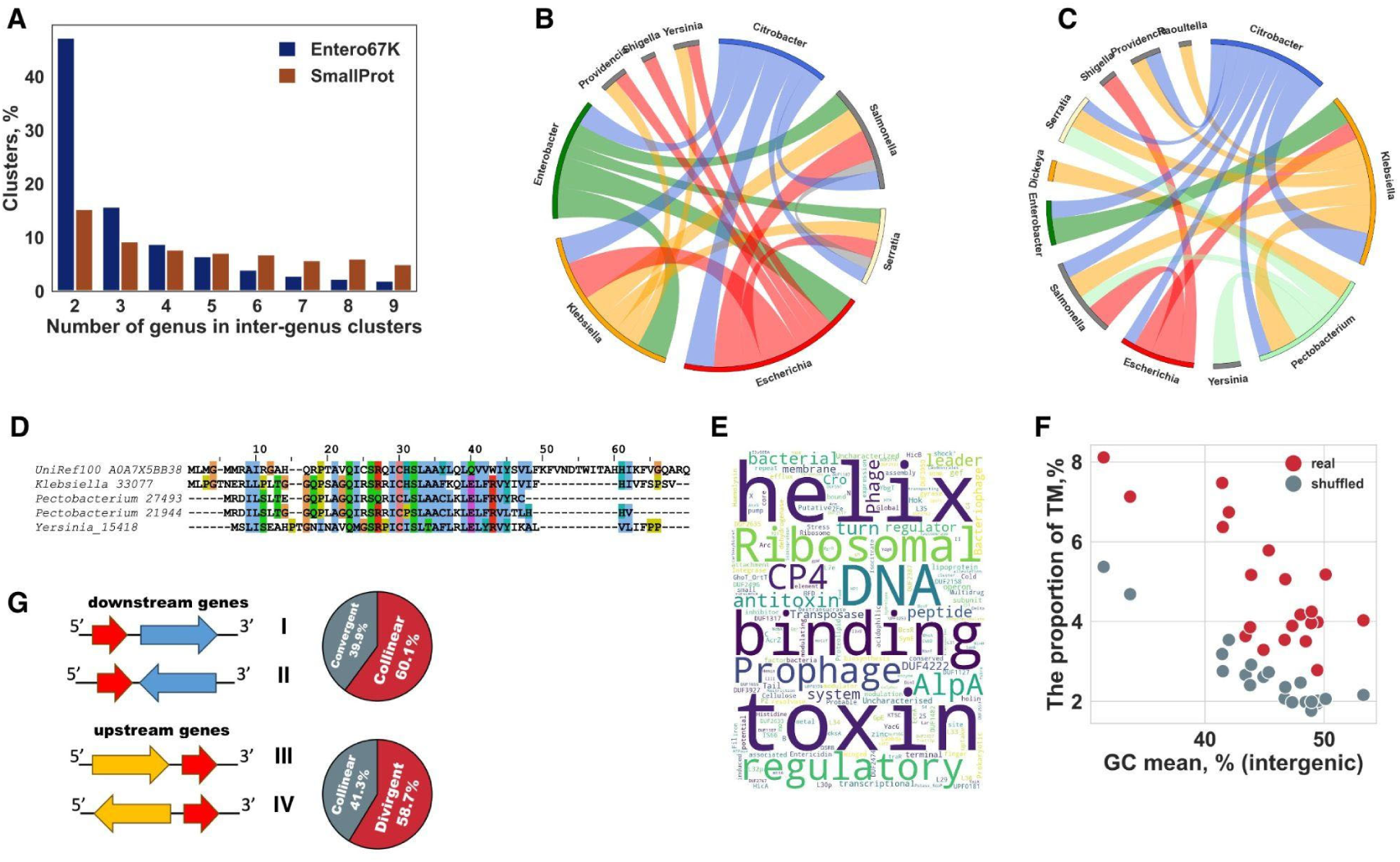
Analysis of the Entero67K families of predicted microproteins. **(A)** The diagram showing conservation of the “MicroProt 6K” families in comparison to small protein families (SmallProt); **(B)** Chord diagram showing top20 interconnections between different genera based on similarity between small protein clusters (SmallProt) revealed by HHsuite; **(C)** Chord diagram showing top20 interconnections between different genera based on similarity between microprotein clusters (Entero67K) revealed by HHsuite; **(D)** Multiple sequence alignments between the Uniref profile representative from Vibrio sp. (UniRef100_A0A7X5BB38) and microprotein clusters from *Klebsiella*, and *Pectobacterium*; **(E)** Small proteins (SmallProt) annotation word cloud. Protein domain names were obtained from the “Pfam” database; **(F)** The scatterplot shows the mean proportion of predicted (TMHMM 2.0^54^) transmembrane domains in Entero67K (“real”) and “shuffled” microproteins (predicted on shuffled intergenic sequences) for each genus and the GC content of intergenic sequences across *Enterobacteriaceae* genera. Entero67K microproteins are shown in red. Genus *Buchnera* and *Candidatus* spp. have the lowest GC content of IGs and the highest percent of “shuffled” microproteins with predicted TM domains; **(G)** The orientation and location of ismORFs from the “Entero 67K” set in relation to nearby genes; collinear (I, III), convergent (II) and divergent (IV) orientations.

We also searched the putative microprotein sequences from Entero67K against the UniProt Reference Clusters database (UniRef30; contains about 260 million protein profiles) and identified 15% (10,076) of Entero67K clusters that matched 11,713 records from UniRef30. About 57% of these Entero67K clusters matched UniRef records from the family *Enterobactariaceae*, such as *Photorhabdus*, *Lelliottia*, and *Xenorhabdus* (Table S5). Homologs in more distant species were also identified. As an illustration, several Entero67K clusters showed significant similarity to the DUF3265 domain-containing protein from *Vibrio* sp. V28_P6S34P95 (UniRef100_A0A7X5BB38 cluster) (Figure 4D). Thus, microproteins, due to their small size, appear to quickly enter the “twilight zone”, where sequence similarity is no longer detectable.

Altogether, the median number of widely-conserved Entero67K microproteins was 83.5 ismORFs/genome (IQR = 36.6–170.25; ∼14% of all Entero67K ismORFs per genome). Thus, a substantial minority of the predicted microproteins were conserved over relatively large evolutionary distances and, by implication, were likely to perform biologically important functions.

#### Most microprotein families have unknown functions

InterProscan 5.60–92.0^46^ was used to annotate the microproteins and small proteins. As could be expected given the strategy employed for microprotein prediction, 99.9% of the Entero67K clusters were not assigned any protein domains from the Pfam database. In contrast, ∼67% of the annotated small proteins in the SmallProt dataset were assigned to known domains (Table S1). The majority of the annotated small proteins were diverse toxin and antitoxins, and ribosomal proteins (see Figure 4E).

The Entero67K microproteins were, on average, significantly more hydrophobic than the annotated small proteins (SmallProt dataset) in all genera (Mann-Whitney U test, *P* < 0.00001 for all comparisons; Figure S2. Related to Figure 4). However, only 5% of the microproteins contained predicted TM domains, compared to ∼15% in SmallProt. However, among the Entero67K microproteins longer than 30 aa, ∼15% contained predicted TM domains, similar to SmallProt, suggesting that the difference between the two sets was due to the smaller size of many microproteins that are too short to be integral membrane proteins.

We further compared the content of predicted TM domains in Entero67K microproteins and random microproteins predicted in shuffled intergenic sequences (k=3, see Methods). In most cases (except for *Buchnera* and *Candidatus* spp. that had the highest AT content in intergenic sequences), significantly more TM domains were predicted for the real ismORFs (Fisher’s exact test, *P*<0.00001, Figure 4F). These findings indicate that, although the high hydrophobicity of the predicted microproteins is a simple result of the high AT-content of the IGs, the formation of transmembrane domains could be affected by selection.

To identify microproteins that could be secreted and involved in cell-to-cell communications, we predicted signal peptides using two methods, Phobius^47^ and SignalP 6.0^48^. A cluster was classified as a secreted microprotein if ≥80% of the sequences were predicted to contain a signal peptide. Overall, we identified 2,168 Entero67K clusters that were predicted to be secreted by at least one of the tools (Table S2).

Thus, our analysis identified a previously unrecognized set of putative small hydrophobic microproteins that appear to evolve under selection. Thanks to the AT-richness of the intergenic regions, hydrophobic microproteins are particularly likely to evolve *de novo*.

#### Genomic context of ismORFs

We next examined the genes adjacent to the Entero67K ismORFs that potentially might belong to the same operons as the ismORFs. We found that ismORFs often co-occurred with genes annotated as “hypothetical”, “transporters” or “transcriptional regulators”. The “hypothetical” proteins and microproteins might represent arrays of functionally interacting yet uncharacterized genes^49,50^.

Although intergenic regions in bacteria are relatively short, the mutual orientation of ismORFs and the adjacent genes could shed light on the mechanism(s) of the emergence of functional microproteins. About 60.1% of the Entero67K ismORFs were co-directed with the downstream genes, which is significantly more than expected by chance (Figure 4G; Fisher exact test, *P* < 0.00001), and 28% of the ismORFs were co-directed with both the upstream and downstream genes (“head-to-tail” arrays). Conversely, the fraction of the ismORFs in the divergent orientation (”head-to-head”) with the corresponding upstream genes (58.7%) was significantly higher than expected by chance (Figure 4G; Fisher exact test, *P* < 0.00001). Conceivably, these preferential orientations facilitate the expression of the ismORFs and thus the emergence of microproteins^22,51^.

It has been shown previously that sequences located downstream of stop codons of many protein-coding genes are subject to selection because of read-through translation^52^. We hypothesized that such read-through events could be involved in the *de novo* birth of functional ismORFs. Indeed, we found that “collinearly” oriented ismORFs with Ribo-seq signals were significantly enriched downstream of protein-coding ORFs (Z test, *P* < 10^-10^) in all examined datasets. Thus, regions downstream of protein-coding genes might be a “hotspot” for *de novo* gene emergence. The majority of these ismORFs might be “proto-genes,” as was previously shown in eukaryotes^4^.

About 13% of the Entero67K ismORFs co-directed with downstream genes had stop codons within 10 nucleotides of the downstream ORF start. Approximately 37% of these ismORFs, overlapped the respective downstream genes by 1–4 nucleotides. Similarly, about 13% (∼35% interect upstream gene) of the ismORFs co-directed with upstream genes started within 10 nt of the stop codon of the adjacent gene. Thus, up to 27% of potentially coding ismORFs seem to emerge in close proximity to upstream and downstream protein-coding genes and can be predicted to be recruited into the same operons with these adjacent genes^53^.

### Microprotein structure prediction

#### Most bacterial microproteins are predicted to adopt simple structures

We sought to predict the structures of the putative bacterial microproteins using the powerful AlphaFold2 (AF2) tool^55^. To reduce the computational time, we first clustered all predicted microproteins from Entero67K at 50% sequence identity; about 88% of the resulting clusters were genus-specific, confirming the low evolutionary conservation of the microproteins (Figure 5A). Next, AF2^56^ was used to predict the structures of cluster representatives. The 11,841 clusters (median = 7 members; “Structured ismORFs”; Table S6) with confidently predicted structures (mean pLDDT scores >80; all 64,896 structures with pLDDT > 60 are available on Zenodo) were analyzed further. As could have been expected given the small size of the microproteins, the majority of the predicted structures contained few secondary structure elements, frequently, only a single alpha-helix or a beta or alpha hairpin (Figure 5B). The N-terminal and/or C-terminal regions were typically unstructured, likely intrinsically disordered^57,58^; the structures of many other microproteins appeared to be completely disordered, without a reliable structure prediction^59–61^. Indeed, we found that, compared to the entire Entero67K, the fraction of microproteins containing intrinsically disordered regions predicted by InterProscan 5.60–92.0 (MobiDB-lite) was substantially greater in the subset with pLDDT score <80 (Z test, *P* < 10^-10^).

**Figure 5.**
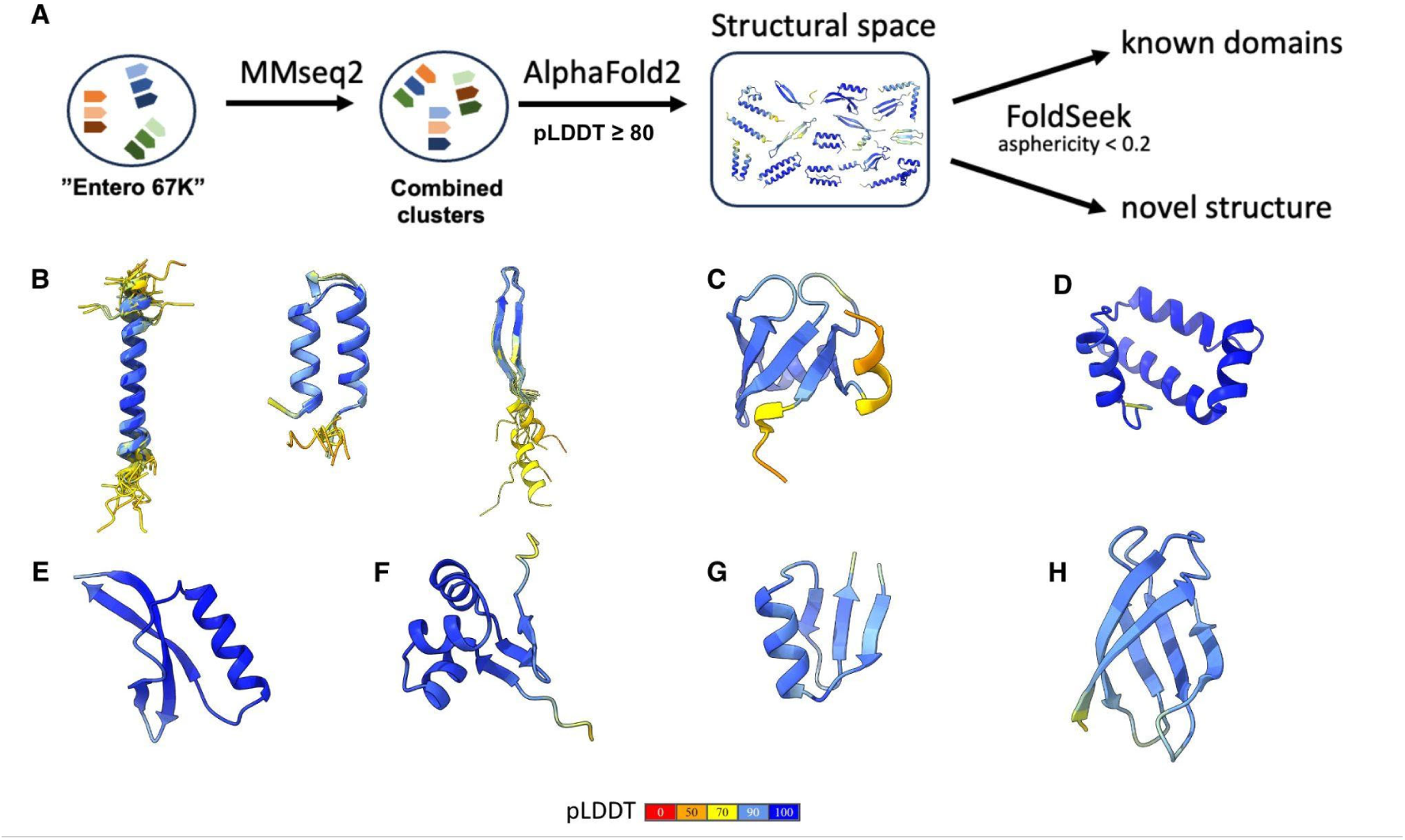
Prediction of microprotein structures using AlphaFold2. **(A)** AlphaFold2 was used to predict the structures of microproteins. Structures with pLDDT > 80 were selected for further analysis. The predicted structures were searched against PDB database; **(B)** Examples of the most frequent secondary elements and structural motifs identified in a set of microproteins; **(C)** An example of a microprotein structure (NC_013353.1_ORF.61612) that resembles the SH3 domain with a beta-barrel-like fold. This structure has hits from the PDB database (for example, 2cud_A); **(D, E, F, G, H)** Examples of structures that cannot be identified in the PDB database from clusters NC_018522.1_ORF.60593, NZ_CP043332.1_ORF.30507, NZ_CP011254.1_ORF.115569, NC_015663.1_ORF.39606, and CP056474.1_ORF.21448, respectively.

Alpha-helices accounted for about 78% of all residues in “Structured ismORFs”, with the highest proportion found in the genera *Buchnera*, *Proteus*, and *Providencia* that have genomes with high AT-content. Apparently, AT-rich IGs encode putative microproteins with biased amino acid compositions that cannot form complex secondary structures. Moreover, 90% of the microproteins were predicted to adopt all-alpha folds (>40% of amino acids in alpha-helices), such as alpha-helix hairpins, bundles, and helix-turn-helix domains. These findings are consistent with previous observations of a positive correlation between the proportion of alpha-helices and pLDDT scores for random and *de novo* proteins^62^.

We also calculated the asphericity of predicted microprotein structures to identify putative globular proteins^63,64^. Most of the putative microproteins are predicted to have elongated structures with asphericity >0.2, which is compatible with the predominance of microproteins containing a single alpha-helix. For further analysis, we selected 107 microprotein structures with asphericity <0.2, to exclude the simplest alpha-helical structures. Foldseek^32^ search yielded 13 (17%) microproteins with 936 significant hits to the PDB database (https://www.rcsb.org; TM score > 0.5; Probability > 0.8). The majority of these hits were proteins with relatively simple folds often implicated in regulatory processes such as transcription regulation, in particular SH3, HTH, or winged HTH domains (Figures 5C and D). We also predicted some distinct structures of microproteins that are not annotated in the PDB database (see examples in Figures 5D, E, F, G H). For example, the cluster of microproteins from Klebsiella (NC_018522.1_ORF.60593) consists of 172 members with four perpendicular helices and a high pLDDT (Figure 5D).

#### Prediction of microprotein dimers

Microproteins might stabilize their structure and increase structural and functional complexity through the formation of homo-oligomers^65^. We evaluated the potential of microproteins for the formation of homo-oligomeric complexes using Alphafold-Multimer (AFM)^56^. Among the predicted microprotein structures with asphericity < 0.5, ∼18% were predicted to form homo-oligomers (with ipTM+pTM > 0.5 and pLDDT > 80; Figure S3. Related to Figure 6). For the SmallProt proteins from *E. coli* UPEC536 (see below), ∼19% (16 of 82) were predicted to form homo-oligomers (Figure S3. Related to Figure 6). Thus, the putative microproteins seem to have a rate of dimerization close to that of annotated small proteins.

**Figure 6.**
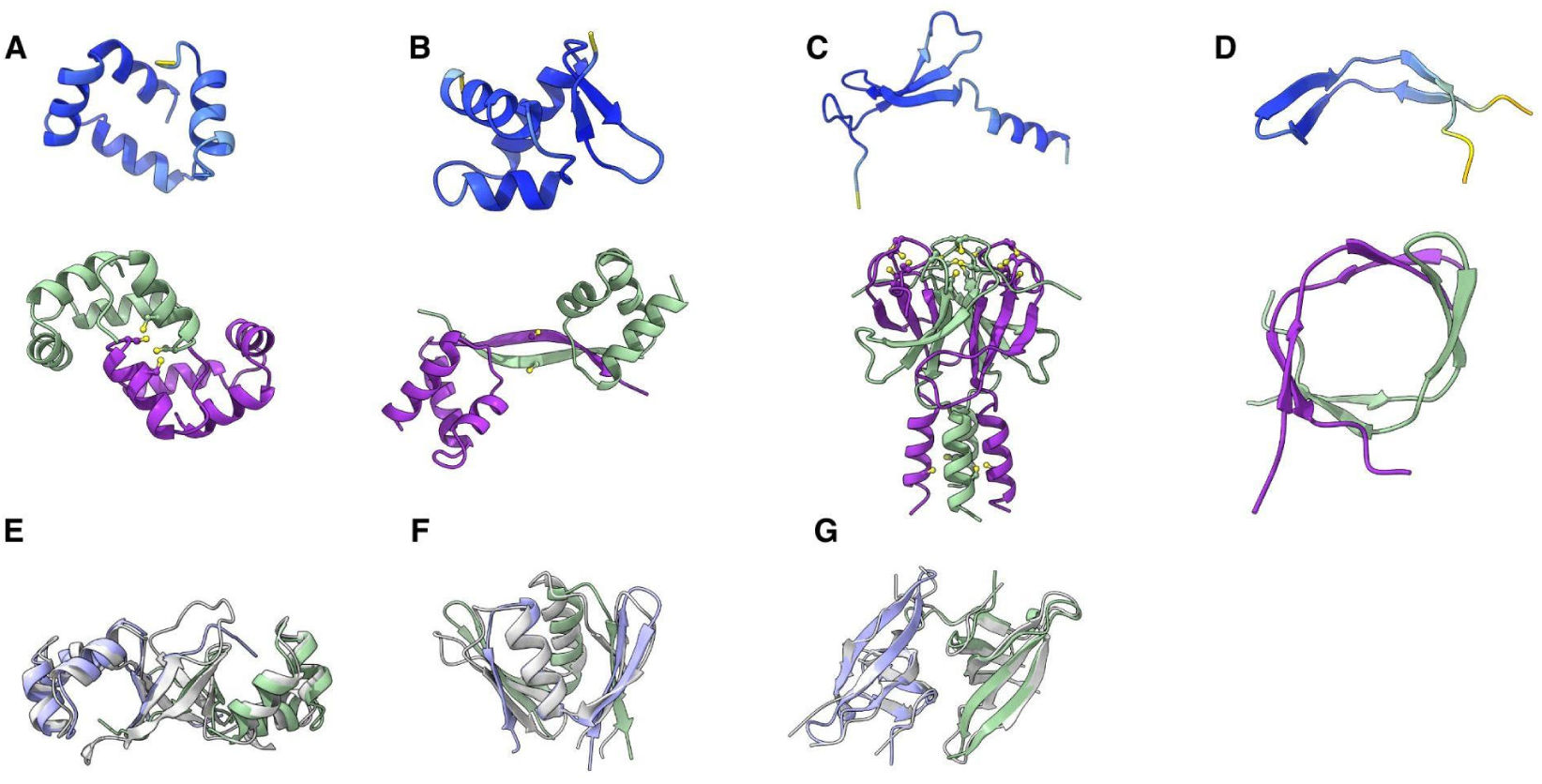
Prediction of microprotein homo-oligomers using AlphaFold2 Multimer. **(A, B, C, D)** Examples of microprotein oligomerization predicted by AFM. Structures with pLDDT > 80 were selected for this analysis. An oligomeric structure is illustrated under each monomer. Protein cartoon structures are colored by pLDDT except in the last row, where homo-oligomers are colored by chain. **(E)** Structure of NZ_CP011254.1_ORF.115569 dimer superimposed with AF-A0A2D5VJZ3; **(F)** Structure of NC_017641.1_ORF.45846 superimposed with DUF3572; **(G)** Dimeric structure of NZ_LR134492.1_ORF.72629 with a beta-barrel-like structure aligned with human SAGA-associated factor 29 using the flexible FATCAT method; The **E**, **F**, **G** chains of the query dimers are colored green and purple, and the target protein is colored gray.

In addition to the dimerization of the simplest folds, such as single alpha-helices or alpha-hairpins, we also predicted different mechanisms of microprotein oligomerizations. For instance, we often noticed cysteines located in close proximity to the predicted dimer interfaces, suggesting disulfide bond formation or metal ion coordination (Figures 6A, B and C), which can be crucial for stabilizing the interaction between chains. Figure 6D illustrates how a beta-hairpin could form a beta-barrel-like structure through dimerization.

Some proteins undergo conformational changes upon dimerization, suggesting that functional activation of these proteins could involve fold switching^66^. An example is a predicted microprotein with a wHTH fold that, in a dimer, is predicted to change the conformation of a beta-hairpin, which unfolds and forms a longer beta-sheet connecting the two HTH domains (Figure 6B). A similar domain-swapping mechanism has been reported previously in a Zalpha-domain-containing protein (ORF112) from Cyprinid Herpes 3^67^.

Although the majority of the predicted interactions in the AFM multimeric models are dimeric, some more complex cases were also observed. For instance, monomers of P072456.1_ORF.26988 are predicted to adopt a simple three-strand beta-meander fold with a putative zinc binding motif and an additional alpha-helix. However, this protein is also predicted to form a tetrameric complex with a core of tightly packed beta-sheets and an alpha-helical bundle (Figure 6C).

The predicted oligomerization of microproteins suggests a mechanism for protein fold evolution whereby the complexity of emerging folds could increase through dimerization, duplication and fusion of *de novo* genes. With this in mind, we represented the predicted microprotein dimers as single chains (see Methods) and used them as queries to search PDB and AFDB with Foldseek^68^. A NZ_CP011254.1_ORF.115569 is an example of a microprotein that, in addition to HTH, has three beta-strains that dimerize, forming a beta-barrel-like structure connecting two HTH domains (Figure 6E). A group of uncharacterized bacterial proteins have a similar structure that is formed by a single chain. For example, the aligned structures AF-A0A2D5VJZ3 with the homo-oligomer of the protein mentioned above have a TM-Score of 0.75, RMSD of 2.96, and 115 aligned residues with a sequence identity of 15%. Another example is NC_017641.1_ORF.45846, which forms a dimer resembling a functionally uncharacterized family of conserved small proteins about 100 amino acids in length, DUF3572 (PF12096) (Figure 6F). Superimposed structures have 83 aligned residues with a TM-score of 0.6, RMSD of 2.7, and sequence identity of 24%.

The beta-barrel-like structure of NZ_LR134492.1_ORF.72629 is predicted to form a dimer that resembles SAGA-associated factor 29 (Sgf29), which consists of two similar domains connected with a linker. The superimposition of these proteins with the flexible FATCAT method gives an alignment with an RMSD of 2.12 A and 109 aligned residues with a sequence identity of 8% (Figures 6G).

### Exploring microprotein-protein interactions in stress response

Because the expression of microproteins can depend on bacterial growth conditions^69^, we analyzed a previously published transcriptome dataset from 32 human bacterial pathogens under 11 stress conditions^70^ to search for potential stress-responsive microproteins. We reanalyzed RNA-seq data for three species from our *Enterobacteriaceae* set and identified 259 differentially regulated ismORFs (DE-ismORFs; FC > 1.5; FDR-adjusted <0.05) in *E. coli* UPEC536, 255 DE-ismORFs from *S. enterica* SL1344 and 269 DE-ismORFs from *Y*. *pseudotuberculosis* YPIII from the Entero67K families. The highest proportion of differentially regulated genes across all conditions, including ismORFs, was observed under hypoxia (Figure 7A; Table S7). Overall, we identified 34 DE-ismORFs that were predicted to be secreted by our pipeline and 68 DE-ismORFs containing predicted transmembrane domains.

**Figure 7.**
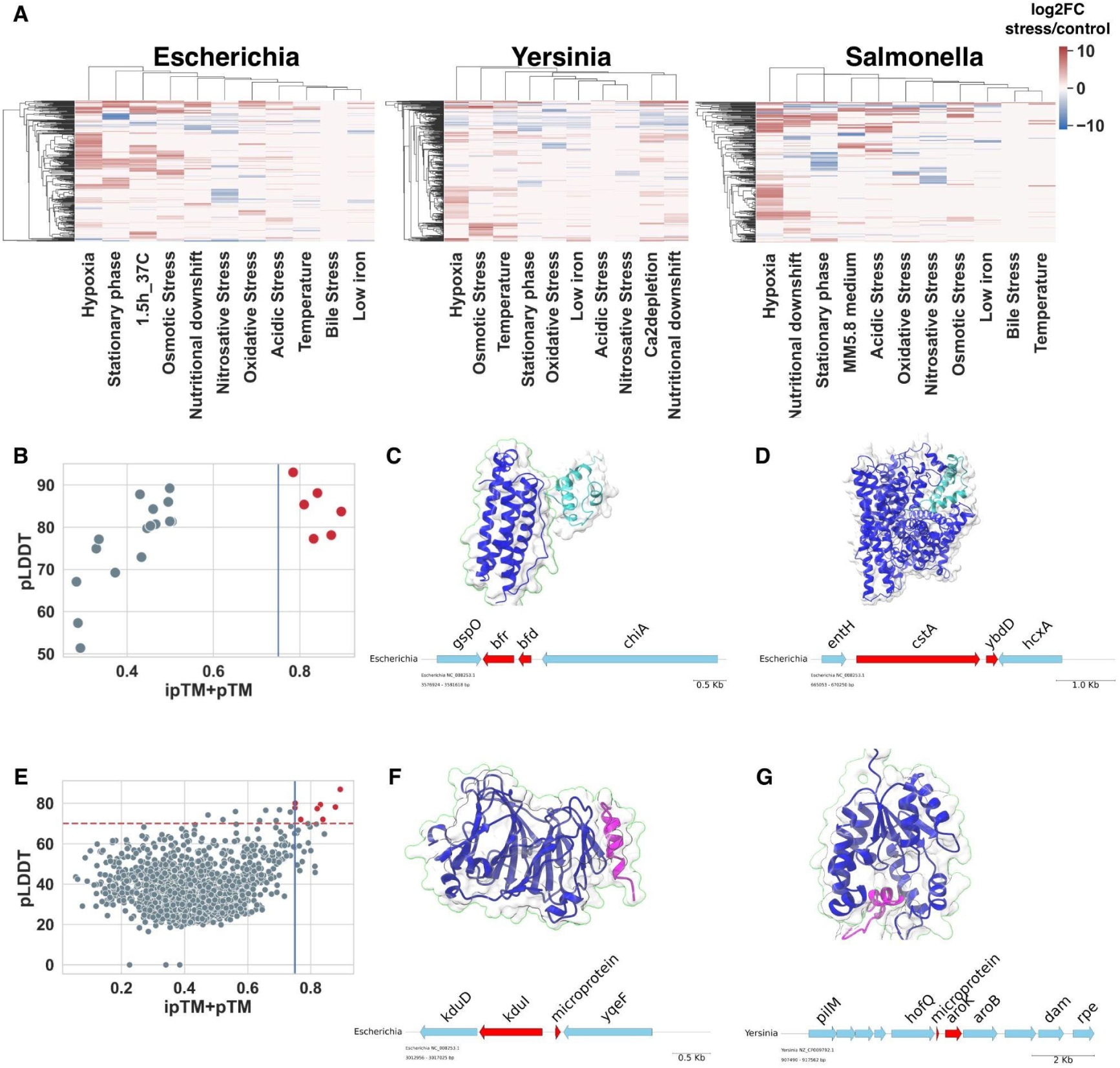
Analysis of protein-protein interactions with AlphaFold Multimer. **(A)** Heatmaps showing transcription regulation patterns of DE-smORF; **(B)** Scatterplot showing pLDDT and ipTM+pTM for experimentally confirmed annotated small protein-protein pair; the vertical line denotes the ipTM+pTM threshold eqtol0.75; 0.75 **(C)** Complex of the small protein Bfd and bacterioferritin Bfr. The interaction of these proteins was confirmed experimentally^76^; the microprotein is shown in cyan; **(D)** Complex of the small protein YbdD and the inner membrane protein CstA; the microprotein is shown in cyan; **(E)** Scatterplot showing pLDDT and ipTM+pTM for microprotein-protein pairs in *E. coli* UPEC536; the vertical and horizontal lines denote thresholds; **(F)** Complex of the microprotein NC_008253.1_ORF.30477 and the protein KduI; The microprotein is shown in magenta; The microprotein is shown in magenta; **(G)** Complex of the microprotein NZ_CP009792.1_ORF.22445 and shikimate kinase AroK; the microprotein is shown in magenta. All protein complexes presented above are predicted with AFM.

Given their small size and lack of known protein domains, the cellular roles of the majority of microproteins are likely to involve modulation of the functions of other proteins via protein-protein interactions^26,71,72^. To assess the potential contribution of this mechanism to stress-regulated microprotein functions, we predicted microprotein-protein interactions using AFM. Despite the experimental and computational limitations, successful prediction of interactions between small proteins and known functional proteins using AFM has been recently demonstrated^73,74^. To define thresholds for reliable predictions, we first predicted 22 complexes between experimentally defined pairs of proteins and bacterial small proteins (Table S8). We observed a bimodal distribution of ipTM+pTM scores and defined ipTM+pTM > 0.75 as the cut-off for further analysis. Because native complexes can contain multiple protein subunits, using only small protein-protein pairs for analysis might explain the observed limitations in predicting such complexes in addition to the limitations of AFM.

Given the high computational cost of AFM prediction of all interactions in the bacterial proteome, we first predicted interactions between annotated small proteins (SmallProt) that could interact with proteins encoded by adjacent genes. Proteins from 8 neighboring genes for each of the 90 differently regulated small proteins from *E. coli* UPEC536 were selected to predict complexes. About 1.3% of such putative complexes, including dimers, passed our cutoff (ipTM+pTM > 0.75), including previously experimentally validated pairs such as AppX-CbdA, Bfd-Bfr, and CstA-YbdD (Figures 7B and C; Table S8).

We next predicted the microprotein-protein complexes between differentially-regulated microproteins and adjacent proteins in *E. coli* UPEC536 and *Y*. *pseudotuberculosis* YPIII. Based on the ipTM+pTM and pLDDT values, 10 (0.5%) microprotein-protein complexes met our stringency criteria (Table S8). The predicted interaction between the microprotein NC_008253.1_ORF.30477 and the protein KduI (5-keto 4-deoxyuronate isomerase; ipTM+pTM=0.83) is shown in Figure 7E. This 23-aa putative microprotein containing an intrinsically disordered region is encoded in a Syn-IGS region, and its transcription level is increased in osmotic and acidic stress conditions (Table S7). The KduI protein facilitates the conversion of galacturonate and glucuronate under osmotic stress^75^, which is compatible with the possibility that this microprotein modulates the activity or stability of KduI. Another conserved syntenic microprotein, NZ_CP009792.1_ORF.22445, is predicted to interact with the binding site of the shikimate kinase AroK (Figure 6F), leading us to hypothesize that this microprotein inhibits this kinase.

## Discussion

Recent large-scale studies revealed an unprecedented diversity of protein families in Earth’s metagenomes^11,12,77^. The majority of these families represent “dark matter”^50^, that is, consist of functionally uncharacterized prokaryotic and viral proteins. Moreover, the actual protein diversity, especially the extent of the dark matter, might be much greater than currently appreciated because our ability to detect small protein-coding genes that, in most cases, appear to emerge *de novo* is limited^11,49,78^. The intergenic sequences span 10-15% of prokaryotic genomes and contain non-coding regulatory sequences as well as genes encoding small regulatory RNAs and small proteins, whose importance in bacterial and archaeal biology is being increasingly appreciated^44,79–81^. However, the prokaryotic microproteomes are far from being comprehensively characterized, in large part because of the difficulty of differentiating ismORFs that actually encode microproteins from spurious ones. At present, annotation of protein-coding genes in the sequenced prokaryotic genome is routinely limited to those containing more than 40 codons, which is indeed a prudent approach because annotation of shorter ORFs as protein-coding genes would result in inexcusable pollution of sequence databases.

In this work, we combined multiple lines of evidence in an attempt to comprehensively predict bacterial microproteomes. Given the scale of the problem, we limited the current analysis to 5668 genomes from the well-studied family *Enterobacteriaceae*. Our main approach was to detect signatures of purifying selection affecting putative microproteins and compare the estimated selective pressure to that affecting functionally and/or structurally characterized bacterial small proteins. As the result, we identified a set of about 67,000 predicted enterobacterial microproteins which translates, on average, into nearly 10% of the bacterial proteomes. In those bacteria for which transcriptomes are available, there is evidence of non-negligible transcription of the ismORFs encoding most of the predicted microproteins. In itself, the presence of an ismORF sequence in a transcript(s) cannot be considered strong evidence of the existence of the respective microprotein but shows that there is at least a potential for microprotein translation. Furthermore, we found that many putative microproteins are encoded in configurations suggestive of operonization with the adjacent genes encoding functionally characterized proteins.

The data on the translation of ismORFs extracted from RiboSeq experiments is more ambiguous. Evidence of translation is available for only a small fraction of the ismORFs predicted to encode microproteins based on selection signatures. Conversely, many ismORFs without significant signs of selection appear to be translated. Definitive interpretation of these observations requires much additional experimentation, including systematic efforts on identification of predicted microproteins. One possibility is that the majority of the ismORFs are translated at low levels under normal conditions but are substantially upregulated under stress. Analysis of the available transcriptomes from stress conditions seems to be compatible with this possibility because many ismORF-encompassing transcripts are indeed upregulated under stress. Conversely, translation of ismORF that are not subject to readily detectable selection could yield a pool of microproteins that are not immediately functional but might have potential for functional recruitment followed by evolutionary fixation.

The great majority of the predicted microproteins have no detectable homologs in protein databases, which is compatible with the origin of the microproteins *de novo*, from non-coding sequences. Structural predictions indicate that most of the microproteins are either intrinsically disordered or form simple, mostly alpha-helical folds such as single helices, hairpins, and helical bundles, including HTH domains. This could be expected given the small size of the microproteins and their likely emergence from random non-coding sequences. A more surprising observation is that a small minority of the microproteins are predicted to form more complex structures, such as SH3-like or distinct beta-barrel-like folds. Although comparatively rare, these observations seem to suggest a path for the birth of protein folds from small ORFs that randomly occur in non-coding sequences. We further predicted that the microproteins could form homooligomers. Given the high probability of the formation of short ismORFs, their products can be considered a dynamic reservoir of initially random, non-functional microproteins that occasionally gain functionality, primarily, through interactions with other proteins. The initial phase of this process might involve positive selection, but then, the microproteins that acquired a function become subject to purifying selection. Similar concepts have been previously developed with regard to the evolution of long non-coding RNAs in eukaryotes^82^.

For most of the bacterial microproteins that are predicted to be functional given the selective pressure they are subject to, the specific functions remain to be elucidated. It appears likely that most of them are regulators of the functions of other, larger proteins with which the microproteins form complexes. Several examples of such regulatory roles for very small bacterial proteins have been reported. For example, in *E.coli*, a 15–amino acid SpfP, encoded by Spot42 small RNA, binds the global transcriptional regulator Crp and blocks its ability to activate specific genes, such as the galETKM operon^24^. In *Salmonella,* the 30-aa microprotein MgtR interacts with the Mg^2+^ importer, MgtA, and thereby controls its activity^83^. We predicted some such interactions of microproteins with proteins encoded by adjacent proteins that are likely to belong to the same operon, but comprehensive computational and experimental analysis of such complexes remains to be performed.

This work provides an extensive resource for the study of bacterial microproteomes. The turnover of *de novo* genes in particular species can be measured by utilizing our data on coding potential, synteny analysis, and evolutionary conservation. By intersecting the coordinates of identified microproteins with experimental data, researchers can select specific candidates for further analysis.

## Limitations of the study

The limitations of this work stem primarily from the high computational costs of some of the analyses. In particular, the prediction of microproteomes was limited to the genomes of bacteria in the family *Enterobacteriaceae*. Distinct trends might exist in other prokaryotes, particularly those with substantially different GC contents of the genomic DNA. The analysis of coding potential was based on estimating evolutionary rates that decreased the number of highly conserved microproteins, or small proteins. Complex formation between microproteins and larger proteins, the functions of which could be microprotein-modulated, was predicted only for a limited set of the putative microproteins that were upregulated under stress, and only the possibility of complex formation with proteins encoded by adjacent genes was assessed. However, the AFM pipeline employed here can be used for the assessment of experimental data, such as comprehensive interactome analysis of bacterial small proteins.

## STAR Methods

### Key resources table

| REAGENT or RESOURCE | SOURCE | IDENTIFIER |
| --- | --- | --- |
| **Deposited data** |  |  |
| All original data and code produced in this work | This paper | https://zenodo.org/records/10672687 |
| Original code produced in this study | This paper | https://github.com/IgorFesenko/prokariotic_smORFs |
| **Software and algorithms** |  |  |
| CD-HIT | Fu et. al. | https://github.com/weizhongli/cdhit |
| RNAcode | Washietl et.al., 2011 | https://github.com/ViennaRNA/RNAcode |
| MEGA X | Kumar et. al., 2018 | https://www.megasoftware.net/ |
| MAFFT v7.475 | Katoh and Standley, 2013 | https://mafft.cbrc.jp/alignment/software/ |
| HH-Suite | Steinegger et al., 2019 | https://github.com/soedinglab/hh-suite |
| orfipy | Singh et.al., 2021 | https://github.com/urmi-21/orfipy |
| Biopython | Cock et.al., 2009 | https://biopython.org/ |
| IQ-Tree | Nguyen et al., 2015 | http://www.iqtree.org/ |
| InterProScan 5.60–92.0 | Quevillon et. al., 2005 | https://github.com/ebi-pf-team/interproscan |
| SibeliaZ | Minkin et.al., 2020 | https://github.com/medvedevgroup/SibeliaZ |
| SignalP 6 | Teufel, 2022 | https://github.com/fteufel/signalp-6.0/blob/main/installation_instructions.md |
| NCBI BLAST+ suite | Altschul et al., 1997; Johnson et al., 2008 | https://ftp.ncbi.nlm.nih.gov/blast/executables/blast+/LATEST/ |
| R | The R Project for Statistical Computing | https://cran.r-project.org/ |
| Python | Python Software Foundation | https://www.python.org |
| MMseqs2 | Steinegger and Söding, 2017 | https://github.com/soedinglab/MMseqs2 |
| uShuffle | Jiang et.al., 2008 | https://github.com/guma44/ushuffle |
| trimgalore 0.6.7 | Krueger, 2019 | https://github.com/FelixKrueger/TrimGalore |
| bowtie 2-2.5.1 | Langmead and Salzberg, 2012 | http://bowtie-bio.sourceforge.net/bowtie2/index.shtml |
| HISAT2 2.2.1 | Kim et al., 2019 | http://daehwankimlab.github.io/hisat2/ |
| FeatureCounts | Liao, 2014 | https://subread.sourceforge.net/featureCounts.html |
| HyPhy BUSTED | Murrell et. al., 2015 | https://stevenweaver.github.io/hyphy-site/methods/selection-methods/ |
| bioinfokit | Bedre, 2020 | https://github.com/reneshbedre/bioinfokit |
| AlphaFold v2.3.2 | Jumper et.al., 2021 | https://github.com/google-deepmind/alphafold |
| foldseek | van Kempen et. al., 2023 | https://github.com/steineggerlab/foldseek |
| HullRad v9 | Fleming, 2018 | http://ec2-52-14-70-9.us-east-2.compute.amazonaws.com/Code.html |
| DSSP v3.1.4 | Touw, 2015 | https://swift.cmbi.umcn.nl/gv/dssp/ |
| TM-align | Zhang, 2005 | https://zhanggroup.org/TM-align/ |
| FATCAT | Li et.al., 2020 | https://github.com/GodzikLab/FATCAT-dist |
| ChimeraX | Pettersen et.al., 2021 | https://www.cgl.ucsf.edu/chimerax/ |

### Lead Contact

All data, including the predicted microprotein structures, is available in the Zenodo repository <u>10.5281/zenodo.10655683</u>. Further information and requests for resources and data should be directed to and will be fulfilled by the lead contact, Igor Fesenko.

### Materials Availability

This study did not generate unique reagents. All data used as inputs for the pipeline is listed in the “data acquisition” section. All outputs of analyses are listed in the “data and code availability” section below.

### Methods Details

#### Extraction of intergenic sequences and prediction intergenic ORFs

Intergenic sequences were extracted from all selected genomes (Table S2) based on RefSeq annotation using a custom Python script. Intergenic sequences with lengths ranging from 33 to 1500 nt were selected for further analysis. For each genome, all ORFs were predicted using orfipy^84^ with the following parameters: “--min 45 --strand b --start ATG,GTG,TTG --stop TAA,TAG,TGA”. Intergenic ORFs were selected based on extracted intergenic sequences using bedtools intersect with the following parameters: “-f 0.8 -u”, resulting in a set of 45,700,542 intergenic small ORFs longer than 15 aa. For each genus, intergenic ORFs with hits to annotated proteins were omitted based on TBLASTN analysis if the *E*-value < 0.00001, the percentage of identity was greater than 50%, and the alignment length was greater than 50% of the microprotein length.

Shuffled ORFs were obtained as follows: 25% of intergenic sequences were randomly chosen for each genus and shuffled by uShuffle^85^ with k=3; ORFs were predicted based on shuffled intergenic sequences using orfipy^84^ as described before. This resulted in 8,484,929 shuffled ismORFs for k=3.

#### Clustering of putative microproteins into families

For each genus, microproteins from 15 to 70 aa encoded by intergenic smORFs were clustered using CD-HIT^25^ with the following parameters: -n 2 -c 0.5 -d 200 -M 10000 -l 5 -s 0.95 –aL 0.95 -g 1. This resulted in 947,440 clusters with at least three sequences. As a control, 562,920 annotated proteins ≤ 70 aa from the RefSeq annotation were clustered into 6107 genus-specific clusters using the same parameters. The proteins annotated as “hypothetical” were excluded from this set.

#### Identification of putative microprotein families

At first, multiple sequence alignments were obtained for each microprotein and small protein cluster using MAFFT v7.475^86^. Then, the nucleotide alignments were converted from protein alignments using pal2nal^87^.

We applied RNAcode^32^ to the microprotein clusters containing at least four unique sequences. Of them, 28,499 clusters with a high diversity (sequence identity < 90%) and RNAcode *P*-values of less than 0.05 were considered to be potentially coding.

The second strategy we developed was referred to as “EvolScore”. For this, average evolutionary rates: *dN* (non-synonymous substitutions), *dS* (synonymous substitutions), and p-distances (*Kd*; Kimura two-parametric model, K2P), were calculated for each microprotein and small protein cluster using MEGA-X^36^. To create a negative control, nucleotide sequences in each microprotein cluster were shuffled using uShuffle^85^ with k = 3 and aligned by MAFFT v7.475^86^. The evolutionary rates were also calculated for these shuffled microprotein clusters. Calculating bootstrap values of 1% and 99% quantities for *dN*, *dS* and *Kd* values in shuffled microprotein clusters and small protein clusters (”SmallProt” dataset), we define the following thresholds for selecting potentially coding microproteins: *dN*<0.2, *dS*<0.7, *Kd*<0.2, and *dN*/*dS*<0.5.

The random forest (RF) algorithm was trained on a set consisting of 3017 “SmallProt” clusters as the positive dataset and 3017 randomly chosen shuffled microprotein clusters as the negative dataset, using evolutionary rates as the feature vectors (70/30 - train/test sets). The performance of the model was assessed using Precision, Recall, and Area Under the Receiver Operating Characteristics (AUROC) from Python sklearn 1.2.2 package^88^.

#### Synteny analysis

SibeliaZ^35^ tool, which aligns closely related genomes using de Bruijn graph analysis, was used for the synteny analysis. A k-mer length of 15, which is the developer-recommended value for bacterial datasets, was used for the runs. As we only required the coordinates of the locally collinear blocks and not the entire nucleotide alignments, we ran SibeliaZ with the (-n) option, as suggested by the developer. Intergenic sequences were intersected with locally-collinear blocks (LCBs) using bedtools^89^.

#### Domains and motifs analysis

Representatives from each microprotein and small protein cluster were scanned for domains using InterProscan 5.60–92.0^46^ with default settings. A cluster was classified as a secreted microprotein family if ≥80% of the sequences in the cluster were predicted to contain a signal peptide by SignalP-6.0^48^ or Phobius^47^. To predict transmembrane domains in putative proteins, TMHMM 2.0^54^ and Phobius^47^ were run with default settings.

#### Conservation analysis

We use HHblits^45^ for performing an all-versus-all profile comparison to find possible homologs for each SmallProt or Entero67K cluster. The obtained matches were filtered using the following parameters: E-value < 0.001, Score > 70, hmm model coverage > 50%, query coverage > 80%. In addition, the Entero67K cluster was searched against the 2020 UniRef30 database and filtered with an E-value < 0.001, Score > 70, hmm model coverage > 50%.

#### Evolutionary analysis

We used HyPhy’s BUSTED algorithm^43^ to identify small protein and microproteins with evidence of positive selection. BUSTED (Branch-Site Unrestricted Statistical Test for Episodic Diversification) provides a test for positive selection in at least one site on at least one branch.

The input trees for the tests of positive selection were constructed from nucleotide alignment IQ-TREE 1.6.12^90^. Benjamini-Hochberg^91^ method was used for multiple testing corrections. We considered a P-value less than 0.05 as evidence for positive selection.

#### RNA-seq analysis

Accession numbers for sequencing read samples were downloaded from the Sequence Read Archive (SRA, https://www.ncbi.nlm.nih.gov/sra; Table S3). FASTQ sequencing read files were downloaded using the SRA Toolkit version 3.0.0 tool fasterq-dump and quality controls were carried out by FastQC 0.11.9. Sequencing reads from FASTQ files were aligned to the bacterial genomes using HISAT2 version 2.2.1^92^ to produce a SAM file for each sample. SAM files were sorted and converted to binary BAM format using SAMtools version 1.19^93^.

For each ORF, mapped reads were counted using bedtools coverage^89^. An ORF was considered transcribed only if it was fully covered by reads. The expression abundances of mapped reads were counted by the FeatureCounts tool^94^. Normalized raw gene expression counts were converted into Transcript Per Millions (TPMs) values using a Python package bioinfokit^95^. For each ismORF, mapped reads were counted using bedtools coverage^89^. A small ORF was considered transcribed only if its TPM > 1 and it was fully covered by reads. Differential expression analysis was performed by the EdgeR package^96^. The ismORFs were defined as differentially expressed ismORFs (DE-ismORFs) with an adjusted p-value ≤ 0.05 and a fold change > 1.5.

#### RIBO-seq analysis

We downloaded *Escherichia coli* MG1655 and BL21 retapamulin-treated Ribo-Seq (GSE122129) and *E. coli* MG1655 Onco112-treated Ribo-Seq (GSE123675) data from the Gene Expression Omnibus (GEO) repository. The Ribo-Seq ribosome protected fragment data for *E. coli* MG1655 (GSE131514) was downloaded from the GEO repository. The Ribo-Seq ribosome protected fragment data for *E. coli* O157:H7 Sakai (PRJNA395712) were downloaded from the Sequence Read Archive (SRA, NCBI). The Ribo-Seq ribosome protected fragment data for *Salmonella enterica* serovar Typhimurium str. SL1344 (GSE149893) and 14028S (GSE87871) were downloaded from the GEO repository.

Raw reads were trimmed using trimgalore v0.6.7^97^. Reads were mapped uniquely using bowtie 2-2.5.1^98^ with the parameter --very-sensitive-local. Reads mapped to tRNA and rRNA were discarded. The following genome versions were used for mapping: *E. coli* MG1655 NC_000913.3, *E. coli* O157:H7 Sakai NC_002695.2, *S. enterica* SL1344 NC_016810.1, and *S. enterica* 14028S NC_003197.2. For the analysis of TIS and RPF data, the SmORFer tool was used^28^.

#### The structure prediction

Protein sequences were clustered with 50% identity and 80% bidirectional coverage using mmseqs2^99^, and structures were predicted only for representative sequences. We used AlphaFold v2.3.2 with complete databases for building multiple sequence alignments to predict protein structures as well as their homo and heteromeric complexes^100,101^. For single-chain proteins, we predicted five models for each protein, and only the best model was selected for further analysis. For complexes, we predicted five models with three multimeric predictions per model, and the best model was chosen based on the ipTM+pTM score. For predicted structures, we calculated the asphericity with HullRad v9 and secondary structures with DSSP v3.1.4 ^63,64,102^.

We used foldseek^68^ with the 3Di structural alphabet and BLOSUM62-based Smith-Waterman-Gotoh algorithm for alignment to search AlphaFold/UniProt50 v4 and PDB databases using structures of microproteins predicted with AlphaFold as queries. We used TM-align for sequence-independent protein structure alignment, FATCAT for flexible structure alignment, and ChimeraX to visualize and analyze protein structures^103–105^.

#### Statistical analysis

Statistical analyses and visualization were made in Python v.3.9^106^ using modules biopython^107^, scipy 1.7.1^108^, sklearn 1.2.2^88^, seaborn 0.11.2^109^, numpy 1.20.3^110^, pandas 1.3.4^110^. The phylogenetic trees were generated by the Toytree package^111^.

## Acknowledgments

The authors thank Gisela Storz for numerous valuable discussions and critical reading of the manuscript. The authors’ research is supported by intramural funds from the US Department of Health and Human Services (to the National Library of Medicine, EVK and SAS). This research was also supported in part by an appointment to the National Library of Medicine (NLM) National Center for Biotechnology Information (NCBI) Research Participation Program (IF). This program is administered by the Oak Ridge Institute for Science and Education through an interagency agreement between the U.S. Department of Energy (DOE) and the National Library of Medicine (NLM). ORISE is managed by ORAU under DOE contract number DE-SC0014664. All opinions expressed in this paper are the authors’ and do not necessarily reflect the policies and views of NLM, DOE or ORAU/ORISE.

## Author contributions

E.V.K. conceptualized and supervised the project. I.F. and S.A.S. and designed the discovery pipeline. I.F. performed analysis of RNA-seq and Ribo-seq data. H.S. predicted protein structures. S.A.S. performed the evolutionary analyses. I.F. and E.V.K. wrote the manuscript, which was edited and approved by all authors.

## Declaration of interests

The authors declare no competing interests.

## Supplemental information

Figure S1. Venn diagram showing intersection between SmallProt clusters classified as “coding” by RNAcode and EvolScore; ROC curve showing TPR (y-axis) and FPR (x-axis) for the RF model. Related to

Figure S2. The hydrophobicity of annotated small proteins and predicted microproteins from Entero67K across all genera

Figure S3. The prediction of small proteins homo-oligomers using AlphaFold2 Multimer

Table S1.The table with small proteins from SmallProt dataset

Table S2. The table with predicted microproteins from Entero67K dataset

Table S3. Transcribed genes and ismORFs

Table S4. The results of Ribo-seq analysis

Table S5. Detailed information about HHsuite analysis

Table S6. The table describes structural analysis of predicted microproteins

Tables S7. Differentially-regulated genes and ismORFs

Table S8. Detailed information about results of AlphaFold Multimer analysis

## Data and code availability

- This paper analyzes existing, publicly available data. The key resources table lists these dataset accession numbers. Data generated during downstream analysis have been deposited at Zenodo and are publicly available as of the date of publication. DOIs are listed in the key resources table.
- All original code has been deposited at Zenodo and is publicly available as of the date of publication. DOIs are listed in the key resources table.
- Any additional information required to reanalyze the data reported in this paper is available from the lead contact upon request.

## Notes

### Competing Interest Statement

The authors have declared no competing interest.

