## Supplementary figures for "The Cryptic Bacterial Microproteome"

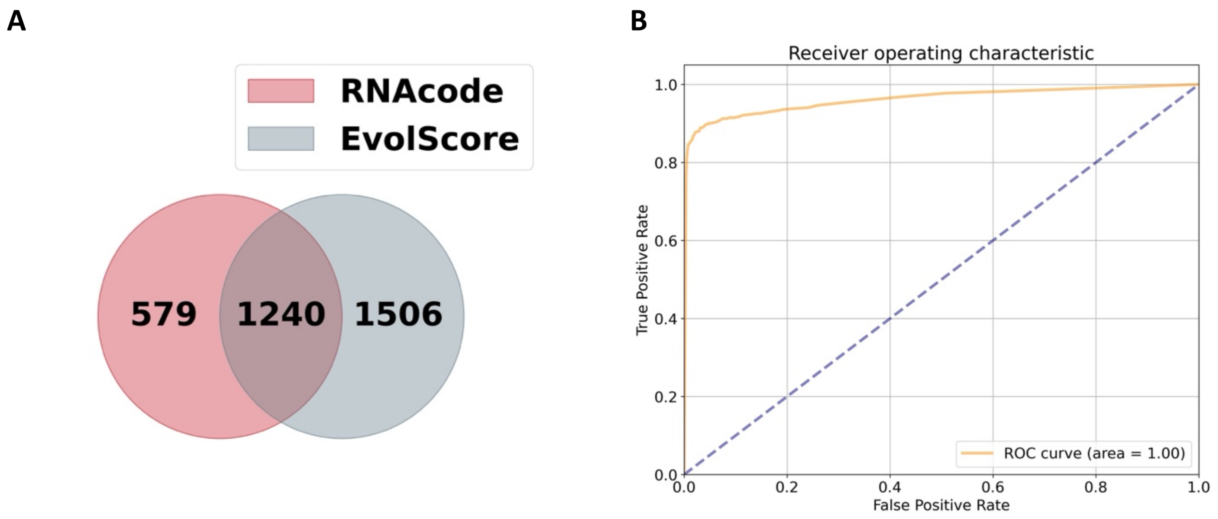


**Figure S1. (A)** Venn diagram showing intersection between SmallProt clusters classified as "coding" by RNAcode and EvolScore; **(B)** ROC curve showing TPR (*y*-axis) and FPR (*x*-axis) for RF model.

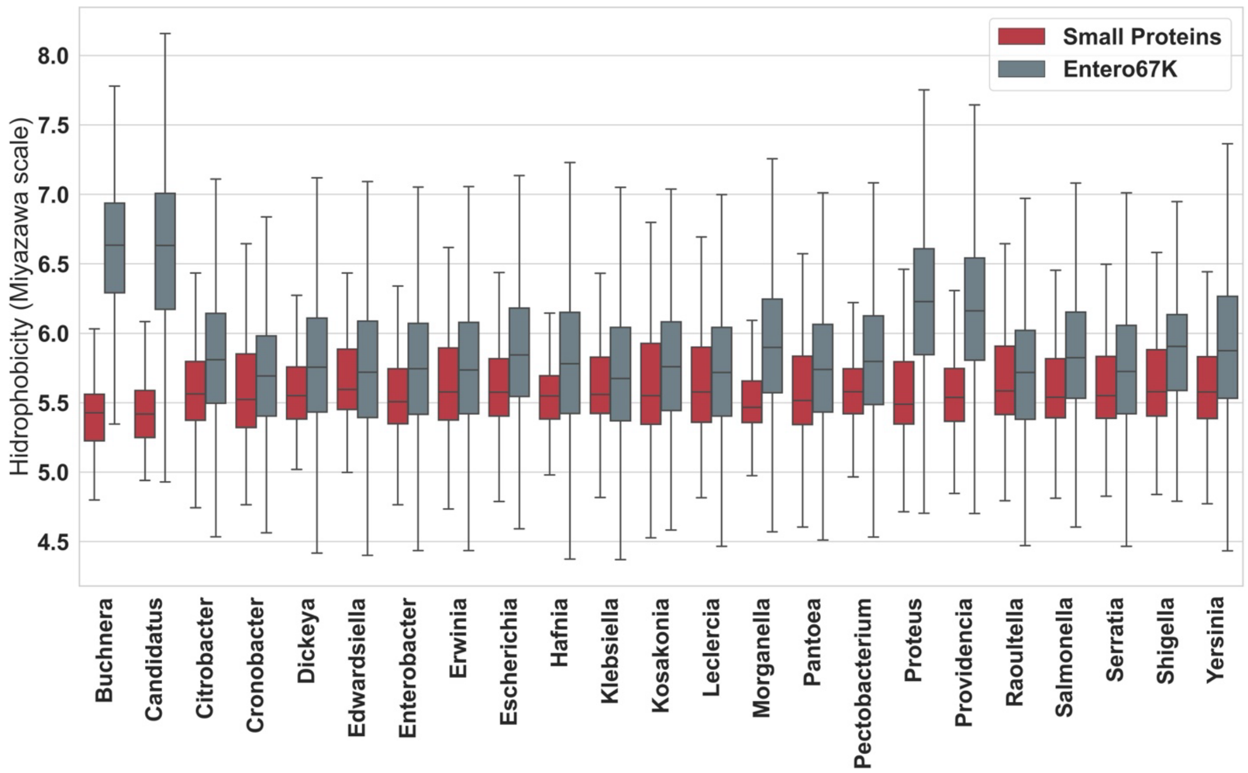


**Figure S2.** The hydrophobicity of annotated small proteins and predicted microproteins from Entero67K across all genera

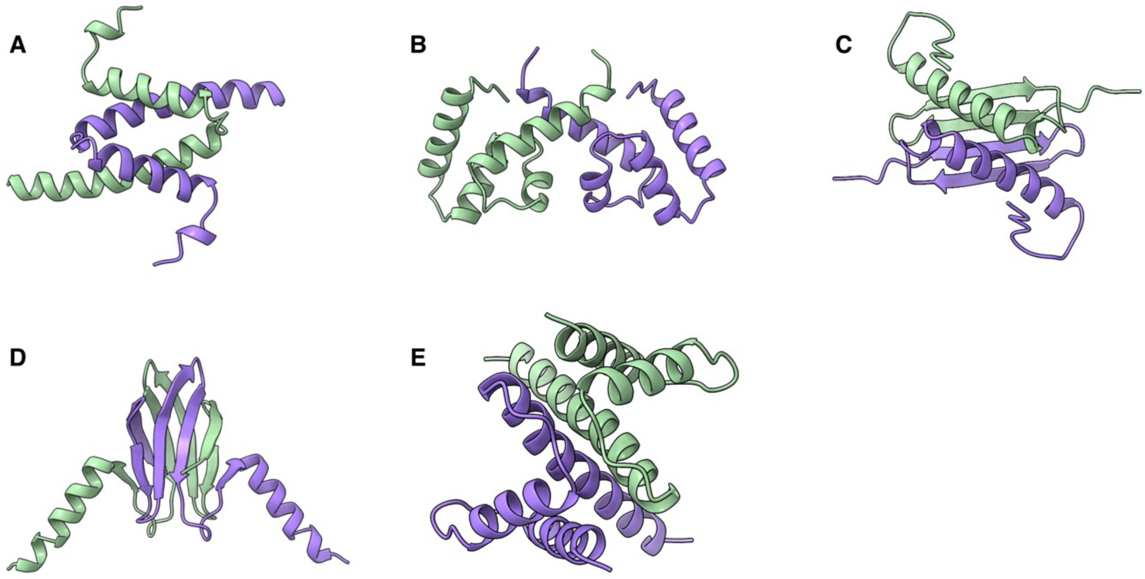


**Figure S3. The prediction of small proteins homo-oligomers using AlphaFold2 Multimer**
**(A)** Structure of RsmS protein dimer; **(B)** Structure of Yjb protein dimer; **(C)** Dimeric structure of YoaG proteins; **(D)** Structure of CsrA protein dimer; **(E)** Structure of FumD protein dimer; The chains of the dimers are colored green and purple.
